# Enhanced JBrowse plugins for epigenomics data visualization

**DOI:** 10.1101/212654

**Authors:** Brigitte T. Hofmeister, Robert J. Schmitz

## Abstract

New sequencing techniques require new visualization strategies, as is the case for epigenomics data such as DNA base modifications, small non-coding RNAs, and histone modifications. We present a set of plugins for the genome browser JBrowse that are targeted for epigenomics visualizations. Specifically, we have focused on visualizing DNA base modifications, small non-coding RNAs, stranded read coverage, and sequence motif density. Additionally, we present several plugins for improved user experience such as configurable, high-quality screenshots. In visualizing epigenomics with traditional genomics data, we see these plugins improving scientific communication and leading to discoveries within the field of epigenomics.

## BACKGROUND

As next-generation sequencing techniques for detecting and quantifying DNA nucleotide variants, histone modifications and RNA transcripts become widely implemented, it is imperative that graphical tools such as genome browsers are able to properly visualize these specialized data sets. Current genome browsers such as UCSC genome browser [1], AnnoJ [2], IGV [3], IGB [4], and JBrowse [5], have limited capability to visualize these data sets effectively, hindering the visualization and potential discoveries with new sequencing technologies. JBrowse is used by numerous scientific resources, such as Phytozome [6], CoGe [7], WormBase [8], and Araport [9], because it is highly customizable and adaptable with modular plugins [5].

Epigenomics is an emerging area of research that generates a significant amount of specialized sequencing data which cannot be efficiently visualized using standard genome browsers. New sequencing technologies such as whole-genome bisulfite sequencing (WGBS) [2, 10], Tet-assisted bisulfite sequencing (TAB-seq) [11], single-molecule real-time sequencing (SMRT) [12], chromatin immunoprecipitation sequencing (ChIP-seq) [13], assay for transposase-accessible chromatin sequencing (ATAC-seq) [14], RNA-seq [15–17], and small RNA-seq [18] have been instrumental in advancing the field of epigenomics. Epigenomic data sets generated from these techniques typically include: DNA base modifications, mRNAs, small RNAs, histone modifications and variants, chromatin accessibility, and DNA sequence motifs. These techniques have allowed researchers to map the epigenomic landscape at high resolution, greatly advancing our understanding of gene regulation. DNA methylation (4-methylcytosine, 4mC; 5-methylcytosine, 5mC; 5- hydroxylmethylcytosine, 5hmC; and 6-methyladenine, 6mA) and small non-coding RNAs (smRNAs) are modifications often found in epigenomic data sets, and function to regulate DNA repair and transcription by localizing additional chromatin marks or inducing post-transcriptional gene regulation [19–21].

We have developed several JBrowse plugins to address the current limitations of visualizing epigenomics data, which include visualizing base modifications and small RNAs as well as stranded-coverage tracks and sequence motif density. Additionally, we have developed several plugins that add features for improved user experience with JBrowse, including high-resolution browser screenshots. These plugins are freely available and can be used together or independently as needed. In visualizing epigenomics with traditional genomics data, we see these plugins improving scientific communication and leading to discoveries within the field of epigenomics.

## IMPLEMENTATION

Plugins are implemented to work with JBrowse’s modular plugin system. Client-side logic, such as visualization, fetching data, and interaction, are written in Javascript relying on the Dojo library [22]. This includes JavaScript classes for viewing data and storing data. Raw data files are standard in genomics, including BAM files for next-generation sequencing reads [23] and BigWig files for quantitative coverage tracks [24]. Python scripts are included to convert output from analysis pipelines to BigWig files needed by JBrowse. Additional styling for each plugin is provided using CSS. Wherever possible, colorblind safe colors were used to improve accessibility.

## RESULTS

### Base modifications

We have developed a plugin to visualize the quantity of 4mC, 5mC, 5hmC, and 6mA at single base-pair resolution. When studying 5mC, the modification is split into two (CG and CH; where H is any nucleotide expect G) sequence contexts for animals or three (CG, CHG, and CHH) sequence contexts for plants, as each context is established and/or maintained by different pathways with different functional roles [20]. Our plugin visualizes the quantity of methylation at each cytosine or adenine using a bar plot (Fig. 1), where values are positive or negative to signify the DNA strand. In most genome browsers, each sequence context must be shown as a different track (Fig. 1a). This is cumbersome when viewing multiple samples and makes it more difficult to determine overlap between context or samples. Our plugin is advantageous because, we color-code 4mC, 5mC, 5hmC, and 6mA sequence contexts and display them on a single track (Fig. 1b, Additional file 1: Fig. S1). However, focusing on a single context or modification can be important, thus our plugin offers several filtering options including by sequence context and base modification.

**Fig 1.**
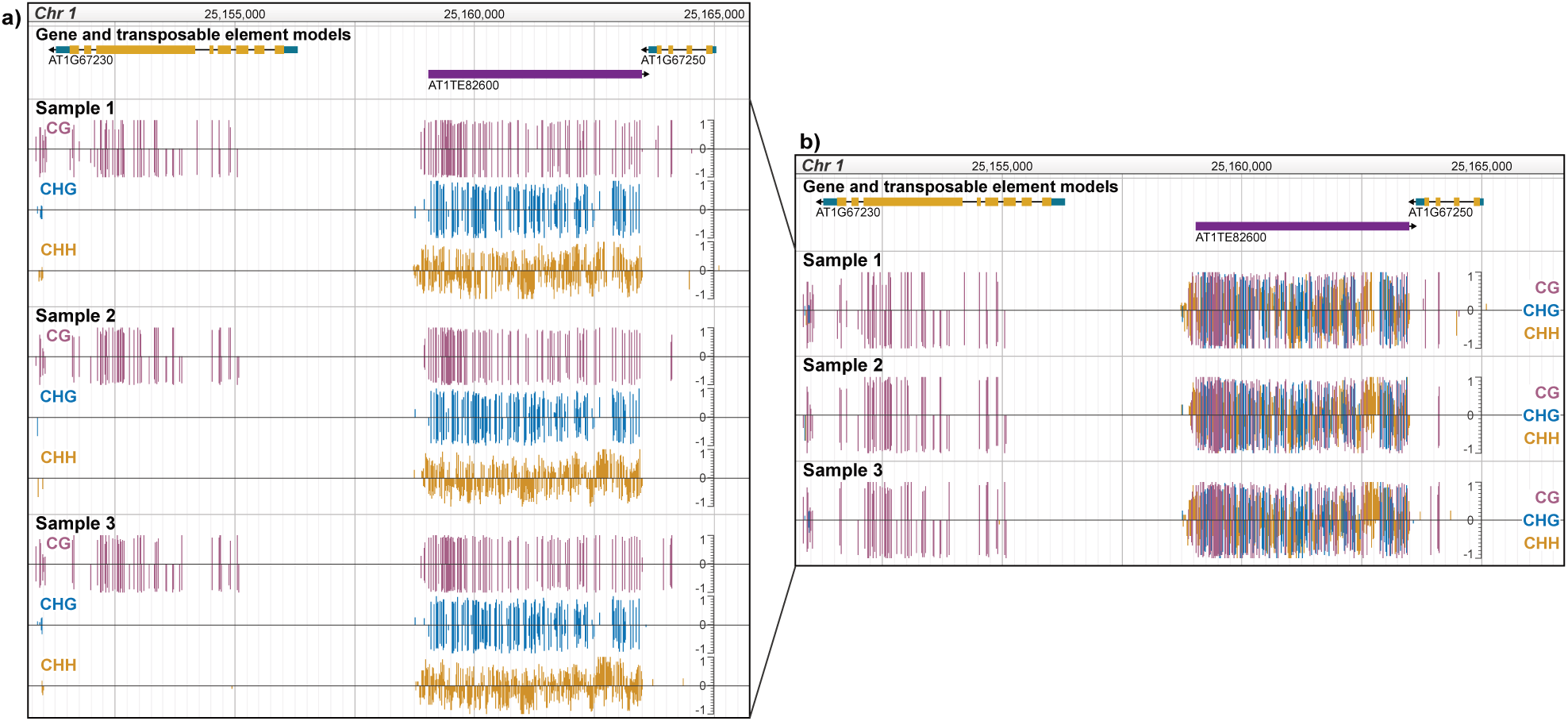
Visualizing DNA base modifications. Top track shows gene models in gold and transposable element models in purple. a) Viewing 5mC in three *A. thaliana* samples without the plugin. b) Viewing 5mC in the same samples with the plugin. For all tracks, height and direction of bar indicates methylation level and strand, respectively. Bars are colored by 5mC sequence context.

### Small RNAs

Currently, JBrowse represents each sequenced RNA as a single read and is colored by sequenced strand (Fig. 2a). When analyzing smRNAs, strand alone does not always provide sufficient information; the size (nucleotides [nt]) of smRNA and strandedness indicate potential function [19]. For example, in plants, 21nt microRNAs can be aligned to single strand and 24nt small interfering RNAs can be aligned to both strands [25]. Products of RNA degradation, however, have varying sizes and align to one strand. To improve smRNA visualization, we color-code reads by smRNA size and retain strand information by placement of smRNAs within the track relative to the y-axis (Fig. 2b). This plugin also includes the ability to filter the reads in a track or multiple tracks by size, strand, and read quality.

**Fig 2.**
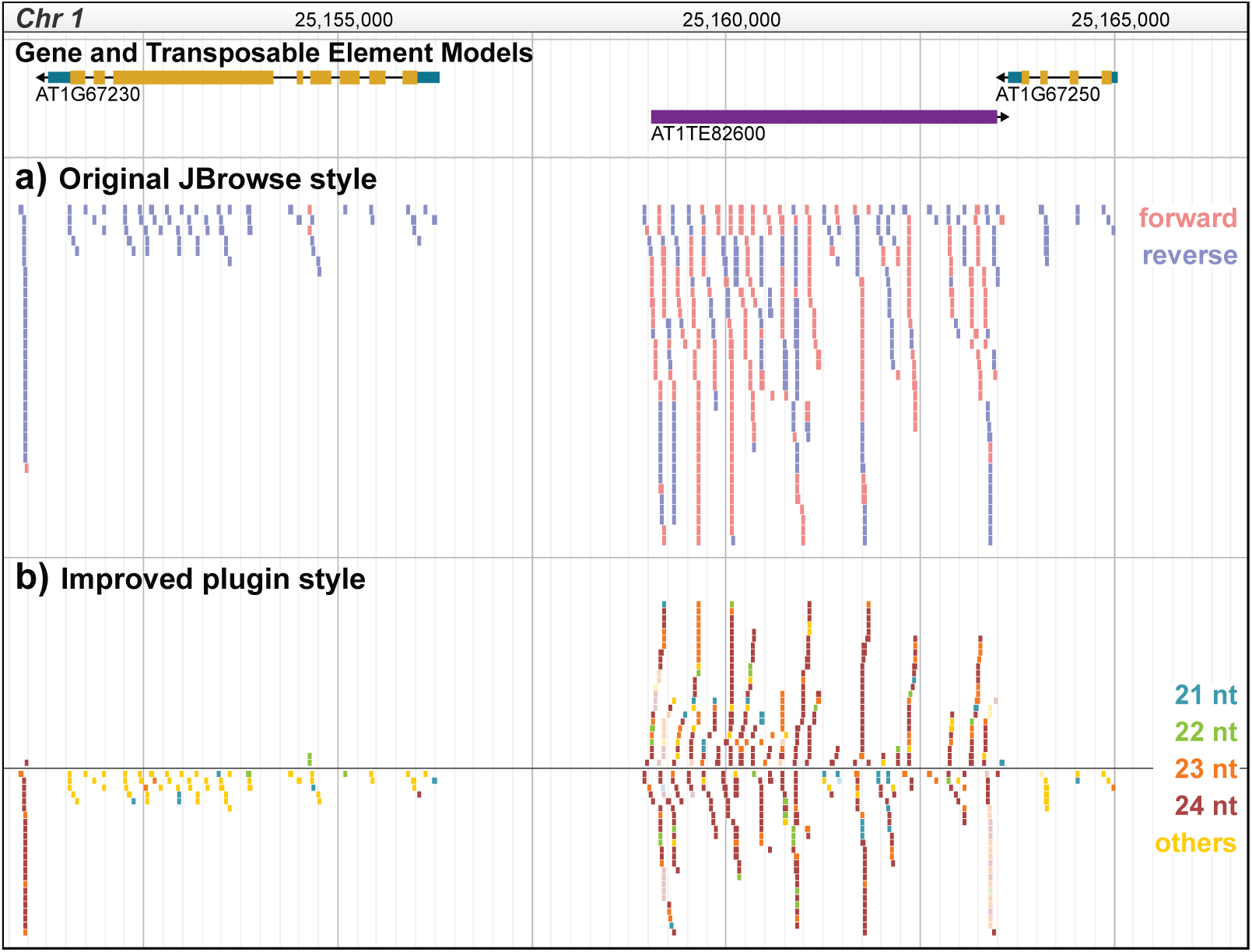
Visualizing small RNAs. Top track shows gene models in gold and transposable element models in purple. a) Viewing smRNA reads, 18 nt - 30 nt, in an *A. thaliana* sample using the general JBrowse alignments track. Color indicates strand; red, forward; blue, reverse. b) Viewing the same smRNA reads using the smRNA alignments track provided by the plugin. Color indicates read length. Position above and below the y-axis origin indicates forward and reverse strand, respectively. Unfilled reads map to multiple genomic locations and filled reads map uniquely.

### Stranded read coverage

Quantitative coverage tracks are necessary for any worthwhile genome browser. It is important for visualizing DNA-protein interactions via ChIP-seq and chromatin accessibility via ATAC-seq where coverage is computed in a strand-independent manner. However, for strand-dependent data types, such as 5mC, small RNAs, and mRNAs, read coverage can greatly vary for opposite strands. The default coverage tracks are unable to handle this, thus we developed a plugin which shows stranded read coverage. For example, WGBS can have uneven coverage on both strands which can make only one strand seem methylated (Fig. 3a).

**Fig 3.**
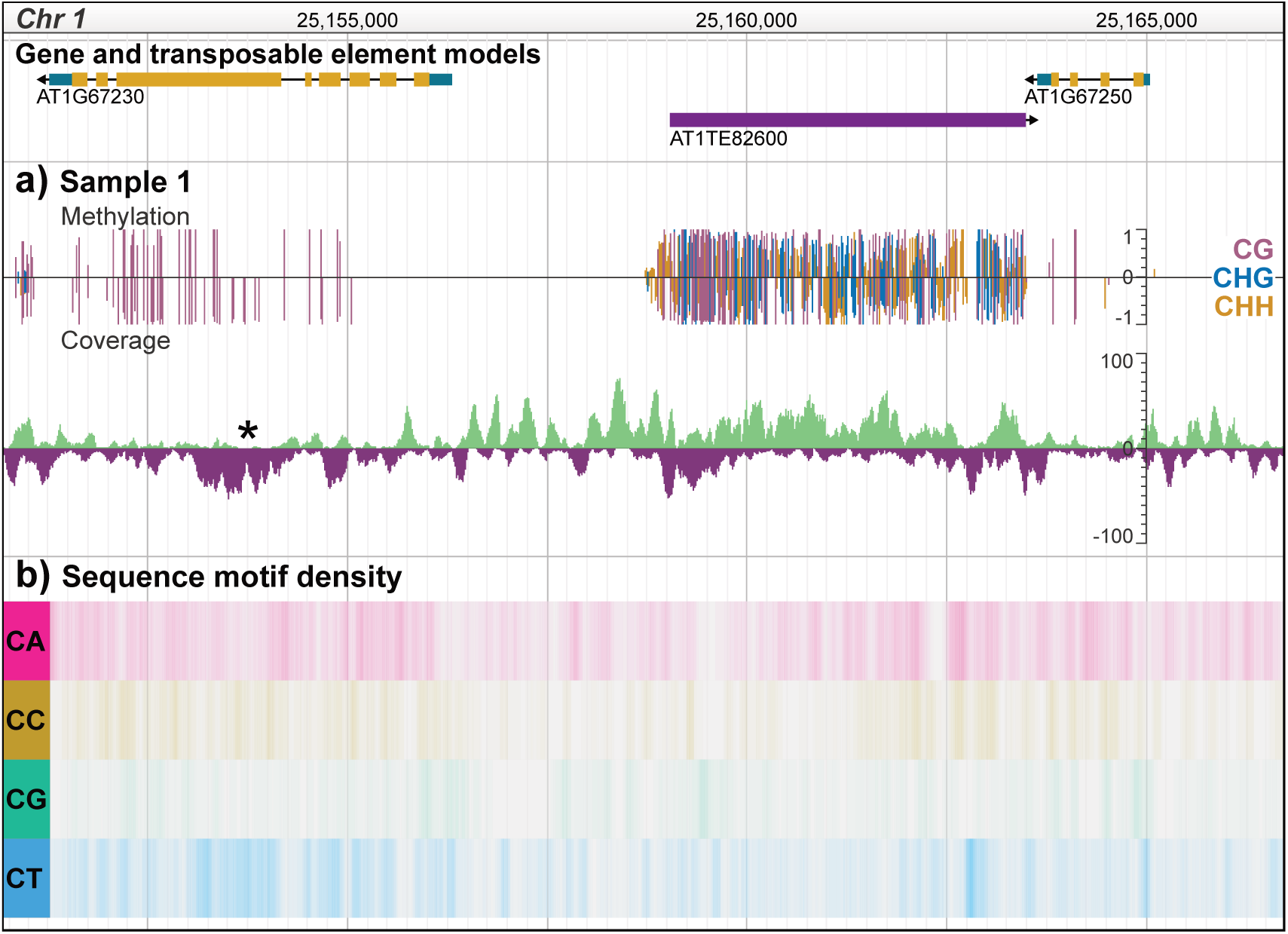
Visualizing stranded coverage and sequence motif density. Top track shows gene models in gold and transposable element models in purple. a) Stranded read coverage for sample used in the methylation track. Asterisk (*) indicates uneven strand coverage which affects the perceived methylation level. b) Dinucleotide sequence motif density in *A. thaliana*. Darker color indicates higher density.

### Motif density

Sequence motifs not only have important roles for protein binding, i.e. binding motifs, but can also impact chromatin formation [26] and recombination hotspots [27]. When correlating the frequency of a sequence motif with another characteristic, i.e. 5mC or histone modification localization, it is preferred to visualize motif density over larger regions compared to single base-pair resolution. To address this, we developed a plugin which visualizes sequence motif density across the genome as a heatmap (Fig. 3b). Users can input multiple motifs in a single track and IUPAC degenerate nucleotides are supported. We also include several options for heatmap coloring and density computation configuration options.

### Exporting browser images

One of the most difficult tasks working with any genome browser is obtaining high-quality screenshots for presentations or publications. We have developed a plugin for JBrowse, which allows the user to take high quality and highly configurable screenshots without installing additional software. A dialog window allows users to set general, output, and track-specific configuration options (Fig. 4). Additionally, our plugin is able to create the screenshot with vector graphic objects, which is preferred for publication-quality screenshots, without needing to change the underlying track configuration parameters.

**Fig. 4.**
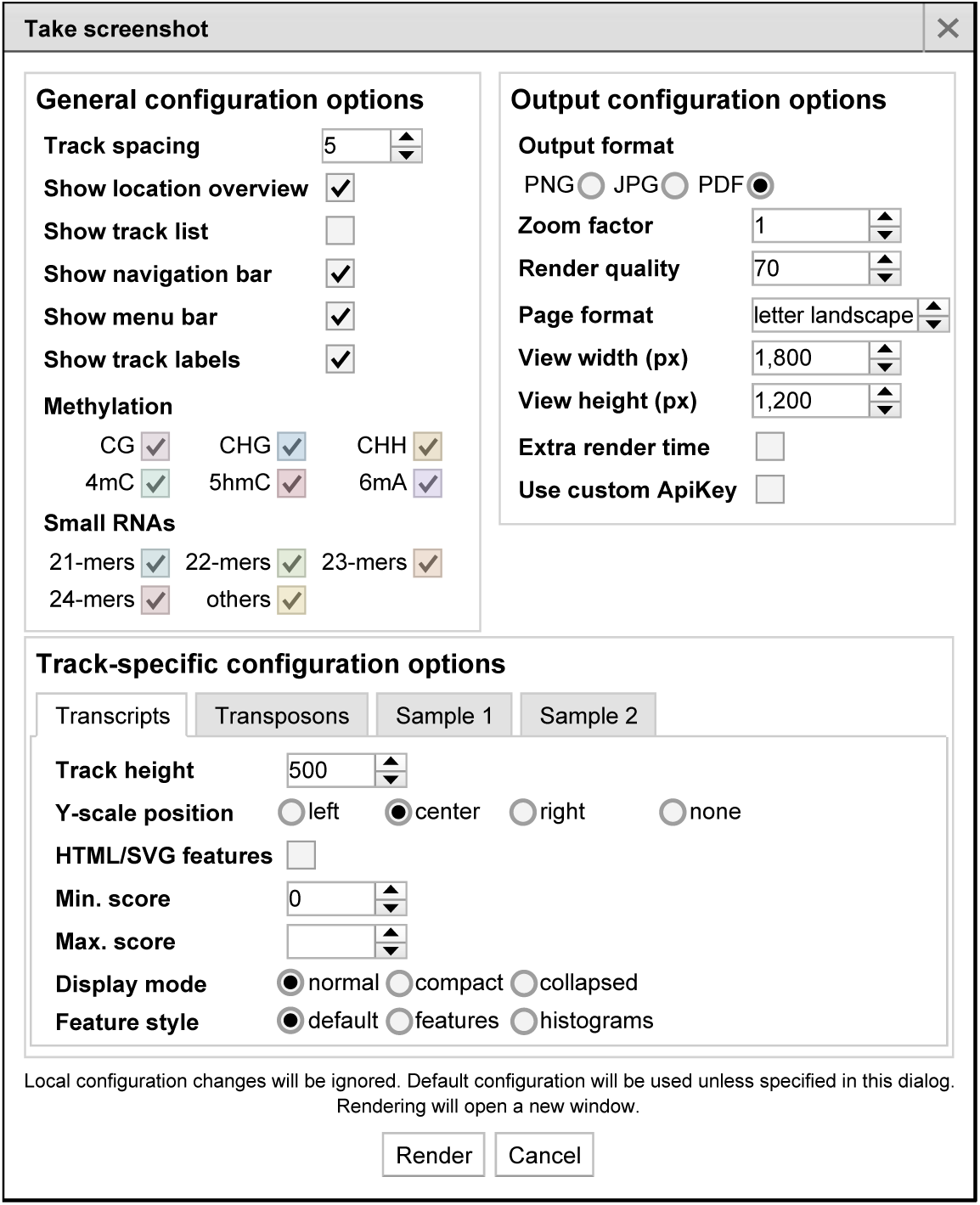
Screenshot dialog window The dialog window that opens when taking screenshots with our plugin. There are numerous configuration options for general visualization, image output, and track-specific settings. This includes exporting each track using vector objects.

### Customization

To improve user experience, we have developed several additional JBrowse plugins. These plugins include: (i) Selecting or deselecting all tracks in a category from a hierarchical track list; (ii) An easily customizable y-scale range and location; and (iii) An option to force a track to stay in “feature” view or “histogram” view regardless of the zoom.

## CONCLUSIONS

With these plugins, we aim to improve epigenomics visualization using JBrowse, a user-friendly genome browser familiar to the research community. All the plugins described can be used together or independently as needed. All plugins are freely available for download and additional customization.

## AVAILABILITY AND REQUIREMENTS

**Project name**: Epigenomics in JBrowse

**Project home page**: http://github.com/bhofmei/bhofmei-jbplugins

**Operating systems(s)**: Platform independent

**Programming language**: Javascript, Python

**Other requirements**: JBrowse 1.11.6+

**License**: Apache License, Version 2.0

**Any restrictions to use by non-academics**: none

## DECLARATIONS

### Ethics approval and consent to participate

Not applicable

### Consent for publication

Not applicable

### Availability of data and material

See Additional file 1 for availability and description of data processing for samples used in the figures.

### Competing interests

Not applicable

### Funding

This work was supported by the National Institute of General Medical Sciences of the National Institutes of Health (T32GM007103) to BTH, the National Science Foundation (IOS-1546867) to RJS., and the Office of Research at the University of Georgia to BTH and RJS.

### Authors’ contributions

Conceptualization and design: BTH and RJS; Implementation and testing: BTH; Writing: BTH; Review and editing: RJS. All authors read and approved the final manuscript.

## Acknowledgements

We would like to thank Adam Bewick, Lexiang Ji, William Jordan, and Melissa Shockey for comments and discussions. We would like to thank Eric Lyons and Colin Diesh for open-source software code that influenced these plugins early in development. We would like to thank all members of the Schmitz lab for using the plugins during development and suggesting additional features. Additionally, we would like to thank Scott Cain and Mathew Lewsey for being early adopters.

## SUPPLEMENTARY FILES

**Additional file 1.** Supplementary figure S1 and supplementary methods. PDF. Additional-file.pdf (103 KB)

